# Effect of protein structure on evolution of cotranslational folding

**DOI:** 10.1101/2020.04.09.033886

**Authors:** V. Zhao, W. M. Jacobs, E. I. Shakhnovich

## Abstract

Cotranslational folding is expected to occur when the folding speed of the nascent chain is faster than the translation speed of the ribosome, but it is difficult to predict which proteins cotranslationally fold. Here, we simulate evolution of model proteins to investigate how native structure influences evolution of cotranslational folding. We developed a model that connects protein folding during and after translation to cellular fitness. Model proteins evolved improved folding speed and stability, with proteins adopting one of two strategies for folding quickly. Low contact order proteins evolve to fold cotranslationally. Such proteins adopt native conformations early on during the translation process, with each subsequently translated residue establishing additional native contacts. On the other hand, high contact order proteins tend not to be stable in their native conformations until the full chain is nearly extruded. We also simulated evolution of slowly translating codons, finding that slowing translation at certain positions enhances cotranslational folding. Finally, we investigated real protein structures using a previously published dataset that identified evolutionarily conserved rare codons in *E. coli* genes and associated such codons with cotranslational folding intermediates. We found that protein substructures preceding conserved rare codons tend to have lower contact orders, in line with our finding that lower contact order proteins are more likely to fold cotranslationally. Our work shows how evolutionary selection pressure can cause proteins with local contact topologies to evolve cotranslational folding.

**Statement of significance:** Substantial evidence exists for proteins folding as they are translated by the ribosome. Here we developed a biologically intuitive evolutionary model to show that avoiding premature protein degradation can be a sufficient evolutionary force to drive evolution of cotranslational folding. Furthermore, we find that whether a protein’s native fold consists of more local or more nonlocal contacts affects whether cotranslational folding evolves. Proteins with local contact topologies are more likely to evolve cotranslational folding through nonsynonymous mutations that strengthen native contacts as well as through synonymous mutations that provide sufficient time for cotranslational folding intermediates to form.

## Introduction

Ribosomes synthesize proteins residue by residue. The ordered emergence of the polypeptide permits cotranslational formation of the protein native structure (1–3). Examples of cotranslational folding processes include forming folding intermediates (4–7), domain-wise protein folding (8–12), and adoption of α-helices and other compact structures in the ribosome exit tunnel (12–14). Cotranslational folding has been shown to enhance protein folding yield by preventing misfolding and aggregation (6, 15–18).

Recent genomic studies from our group and others provide complementary evidence that cotranslational folding has been evolutionarily selected for (19, 20). Specifically, examination of sequence-aligned homologous genes found that rare codons are evolutionarily conserved. Rare codons are translated at slower rates, and translational slowing along the transcript may facilitate formation of native structure (21). The study from our group examined *E. coli* proteins in particular, finding that conserved rare codons are often located downstream of cotranslational folding intermediates that were identified using a native-centric model of cotranslational protein folding (20). In a subsequent computational study using a more realistic all-atom sequence-based potential, we found that the positions of slowly translating, rare codons could correspond to nascent chains lengths that exhibit stable partly folded states as well as fast folding kinetics (22).

While these findings suggest mechanistic reasons for the evolution of cotranslational folding, there is still no clear understanding of which proteins are likely to fold cotranslationally. Many proteins fold posttranslationally (11, 23, 24) or with the assistance of chaperones (25, 26). Additionally, it is unclear to what degree sequences have evolved to optimize either cotranslational or posttranslational folding. To address these questions, we used an evolutionary modeling approach (27). We constructed a fitness function that depends on outcomes of protein translation and folding. We then simulated evolution of coarse-grained lattice proteins, whose folding performance we evaluate using Monte Carlo (MC) simulations of protein translation and folding.

We hypothesized that proteins with local contact topologies would evolve to fold cotranslationally since such topologies are likely to permit piecewise assembly of the native fold. To this end, our evolutionary simulations investigate prototypical lattice proteins of varying contact orders (28). We find that evolved proteins with low contact order fold cotranslationally, with nascent chains adopting native-like conformations beginning when proteins are about two-thirds translated. For high contact order proteins, native structures are only stable toward the end of translation, when translation nears completion. Separately, we assessed the fitness effect of rare codons by simulating translation with a longer elongation interval for individual codons, finding that only proteins that fold early on during translation show improvements in fitness from slow codons at positions aside from the C-terminus. We then performed a bioinformatics investigation using data from our previously published work, which associated rare codons with cotranslational folding intermediates in *E. coli* proteins (20). In agreement with the simulation results, protein substructures preceding rare codons are lower in contact order than random substructures in proteins without conserved rare codons. Our work mechanistically explains how proteins with local contact topologies can evolve cotranslational folding through nonsynonymous mutations that stabilize partial-length native states and synonymous mutations that provide additional time for such native states to form.

## Methods

### Model connecting protein translation and folding to cellular fitness

To simulate evolution of lattice protein sequences, we built a model relating outcomes of protein translation and folding to cellular fitness. A protein undergoing translation can reach its native state during or after translation (Fig. 1A). Following translation, a free protein not in its native state is vulnerable to degradation or aggregation with other proteins, processes assumed here to be irreversible. We consider the survival probability of a protein in the cellular environment over time. Let *S*(*t*) be the probability that a protein survives to time *t* with *t* = 0 being the time the protein leaves the ribosome and *S*(0) = 1. For a protein not in its native state, there is some rate *k_d_* for the protein to be degraded. Based on this model, *S*(*t*) evolves according to the following differential equation:

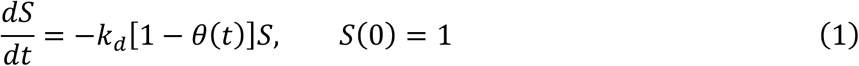

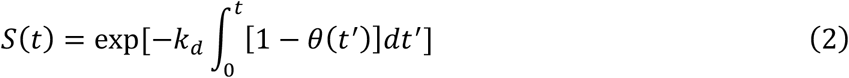

where *θ*(*t*) = {0,1} is an indicator function whose value is 1 when the protein is in its native state at time *t*. Under this model, *S*(*t*) does not decrease if the protein is in its native state. The half-lives of unstable proteins are as short as a few minutes (29–31), compared to hours or days for stable proteins (29, 31, 32), justifying this assumption. A protein thus has a higher survival probability if it folds quickly and remains stably folded.

**Fig. 1:**
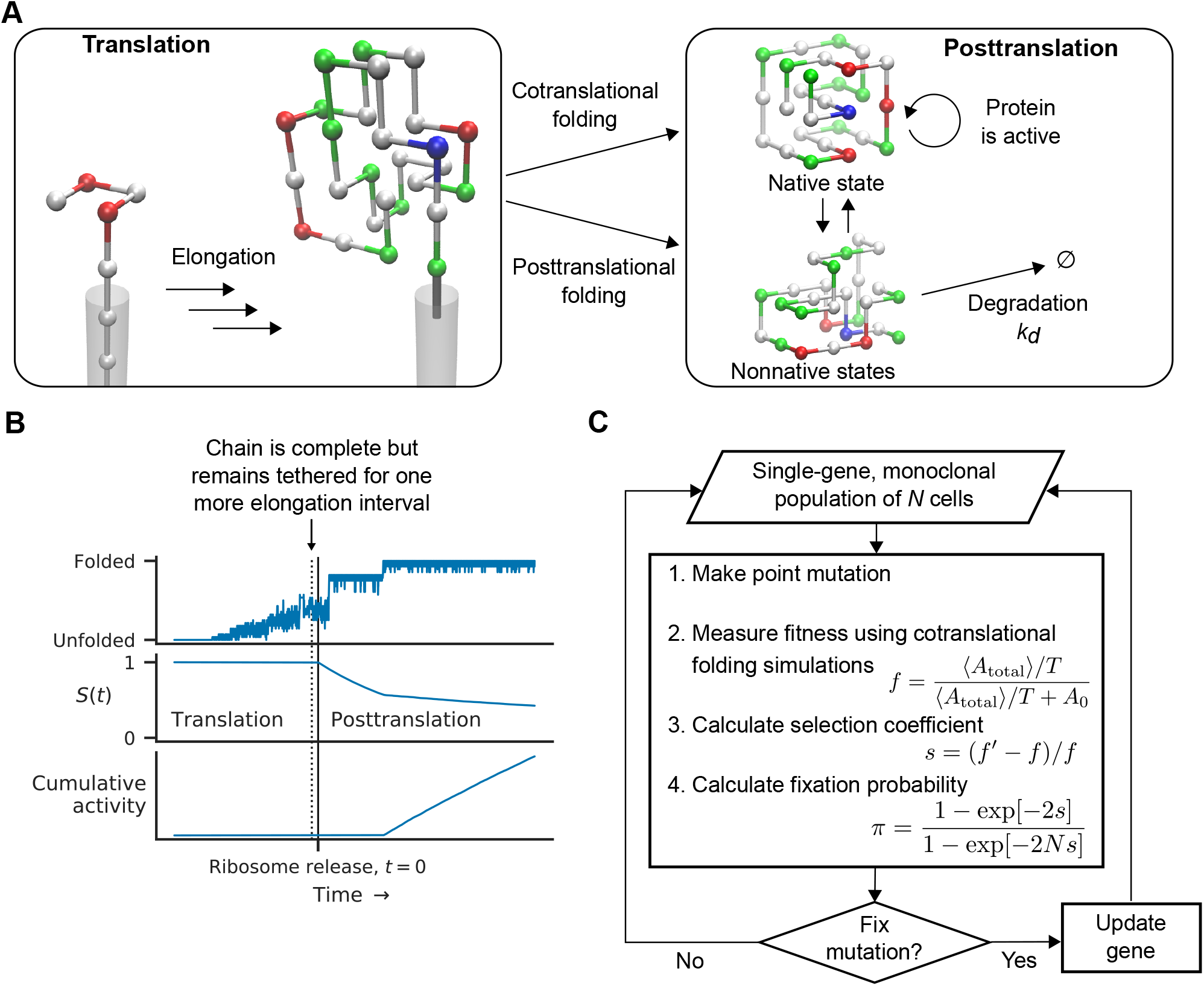
Connecting protein translation and folding to fitness and evolution. (A) Model of protein biogenesis: A protein undergoing translation may reach the native state before or after release from the ribosome. After it is released, the free protein is vulnerable to degradation if it is not in its native state. The protein can only carry out its function when it is in the native state. (B) Example cotranslational folding trajectory in which the protein folds posttranslationally. The point at which the protein is complete but still tethered to the ribosome is indicated by the vertical dotted line. Top: folded/unfolded states depicted by number of native contacts formed. Middle: Survival probability *S*(*t*) (Eq. 2), which decreases after translation if the protein is not in its native state. Bottom: Cumulative protein activity (Eq. 4), which increases when the protein is in the native state. (C) Evolutionary simulation scheme for evolving proteins. For each generation, a trial mutation is assessed via cotranslational folding simulations (as shown in (A)). The time to fold *t*\* and native state stability *P*_nat_ determine total protein activity (Eq. 5), which, averaged over multiple folding trajectories, determines cellular fitness. The mutation is fixed with probability *π*.

We next model how *S*(*t*) affects the activity of the protein over time. We assume a protein can perform its biological function only when in its native state and if it has not been degraded. Based on these assumptions, the total cumulative activity of the protein (e.g. enzymatic output), *A*_total_, is probabilistically described by the following equation:

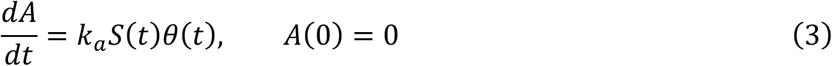

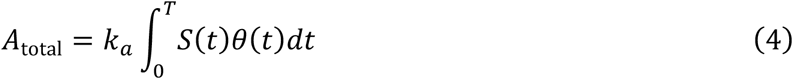

where *k_a_* is an activity rate constant (which we set to 1), and *T* is some long timescale corresponding to the period of time that the protein is biologically relevant, such as the length of the cell cycle.

Fig. 1B illustrates the relationship between protein folding, survival, and activity for a single protein folding trajectory. In this particular trajectory, the protein folds posttranslationally. During translation, the protein is not vulnerable to degradation, and *S*(*t*) is 1. Following release from the ribosome, *S*(*t*) decreases for every time unit the protein is not in the native state. *S*(*t*) decreases at a higher rate during the initial passage to the folded state and then decreases more slowly after the protein enters the native state energy basin and fluctuates in and out of the native state. Protein activity only begins to accumulate after the protein reaches the native state. *A*_total_ corresponds to cumulative activity at a later time *T*.

To reduce the amount of computation required for evaluating the protein activity function (Eq. 4), we assume proteins fluctuate on fast timescales in and out of the native state, occupying the native state with probability *P*_nat_ = 〈*θ*(*t*)〉. After simplifications and applying this fast-fluctuation assumption (see Extended Methods in the Supporting Material), Eq. 4 becomes the following:

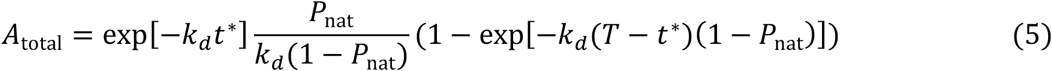

where *t*\* is the first passage time to the native state. According to Eq. 5, *A*_total_ decreases exponentially with *t*\*. On the other hand, the relationship between *P*_nat_ and *A*_total_ is more complex. For realistic values of *k_d_* and with *T* − *t*\* ≈ *T, k_d_T* ≫ 1. If *P*_nat_ is close to 1 such that the entire argument of the exponential is small, *A*_total_ is proportional to *P*_nat_. Otherwise, *A*_total_ is proportional to *P*_nat_/1 − *P*_nat_.

Finally, we relate *A*_total_ to cellular fitness, *f*. For convenience, we divide *A*_total_ by total time *T* to rescale it to the range [0,1]. We treat the protein as essential to cellular growth, and therefore its activity is related to cellular fitness. We use a metabolic flux-type equation to relate total protein activity to *f* (33–36):

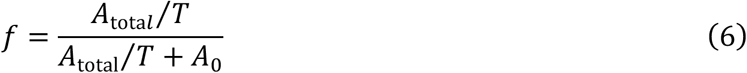

where *A*_0_ is a constant which sets the value of *A*_total_/*T* where fitness is half maximal. Together, Eqs. 5 and 6 formulate how protein folding kinetics and stability determine fitness in our model.

### Evolutionary simulations using a lattice protein model

To explore how proteins evolve under prototypical functional selection, we ran evolutionary simulations that fix or reject mutations in a protein sequence based on the measured fitness. The evolutionary simulation scheme is illustrated in Fig. 1C. We simulate evolution according to a discrete-generation monoclonal model in which, each generation, an arising mutation either fixes (takes over the entire population) or is lost. Our model organism has only a single gene, corresponding to the protein under investigation. Every generation, a mutation is made to the current sequence, and the fitness of the trial sequence, *f*′, is evaluated using protein folding simulations. A selection coefficient is calculated as *s* = (*f*′ − *f*)/*f* (27), and the mutation is fixed with probability

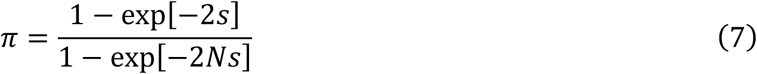

where *N* is the population size. Eq. 7 comes from classical population genetics (37).

In the fitness assessment, we use MC simulations of lattice model proteins undergoing translation. A lattice protein, as illustrated in Fig. 1A, treats protein residues as a connected set of vertices on a cubic lattice. Our model uses 20 amino acid types, whose interaction energy is given by a 20×20 interaction matrix (38).

The MC simulations of translation and folding have two phases: translation and posttranslation. During translation, MC simulation alternates with addition of single residues to the C-terminus of the nascent chain. The length of MC simulation between elongations is set by the chain elongation interval. The nascent chain C-terminus is simulated as if tethered to an extended chain of infinite length (as illustrated in Fig. 1A), representing yet to be translated residues emerging from the ribosome. No lattice protein residues may occupy this space, and there are no interactions between the protein and the untranslated residues. The protein remains tethered for one additional elongation interval after the final residue is added, representing the ribosome release interval (39). During the posttranslation phase, the protein is no longer tethered and has no conformational restrictions. Fig. S2 in the Supporting Material shows example folding trajectories for sequences studied in this work. Because MC simulations are stochastic, multiple trajectories are used to estimate *t*\* and *P*_nat_, and an average *A*_total_ is used in Eq. 6. Additional details on simulation procedures are described in the Extended Methods in the Supporting Material.

The fitness function, as defined by Eqs. 5 and 6, by selecting for protein activity, effectively includes selection on folding kinetics and stability of proteins undergoing translation. As controls, we also ran the same evolutionary simulations under two alternative evolutionary scenarios in which we changed how we assessed fitness. In the first alternative scenario, we skip lattice protein translation. Instead, fitness assessments use MC simulations that begin with full-length proteins in a fully extended conformation, mimicking *in vitro* refolding and thereby making *in vitro* folding speed a determinant of fitness. The second alternative scenario ignores first passage time to the native state. In this case, MC simulations begin with full-length proteins already in their native conformations. *t*\* in Eq. 5 is set to 0, and MC simulations only measure *P*_nat_. Fitness in this scenario therefore depends on stability and does not depend on folding rate, although there may be selection for a slow unfolding rate. We refer to sequences evolved without translation as “evolved, no translation,” and we refer to sequences starting in the native state as “evolved, no folding.” The degradation rate, *k_d_*, and other simulation parameters remained unchanged for these alternative evolutionary scenarios.

### Simulation parameters and analysis

The simulation time unit, *t*, is defined as *t* = MC step/protein length, which accounts for the local nature of the MC move set. The key simulation parameters are the elongation interval and degradation rate *k_d_*. Simulation parameters were chosen so that ratios of timescales between translation, protein folding, and degradation are biologically reasonable. An explanation of how simulation parameters were selected is given in the Extended Methods, and a listing of the parameters is shown in Table S1 in the Supporting Material.

A key characteristic of different lattice proteins simulated in this work is the topology of the native structure that the proteins fold to. Different structures differ in the degree to which residues forming native contacts are separated in primary sequence. Contact order is defined as

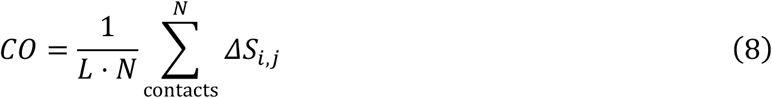

where *L* is the length of the protein, *N* is the number of contacts, and *ΔS_i,j_* is the separation in primary sequence between contacting residues *i* and *j* (28).

Nine lattice protein native structures were selected from the representative 10,000 structure subset of 27-mers used in previous works (40, 41). Each native structure arranges the chain in a 3×3×3 cubic native fold. Three each of low, medium, and high contact order structures were chosen. Initial sequences for evolution were designed to be thermodynamically stable in the selected native conformations via Z-score optimization (42–44); all initial sequence Z-scores were below −50. Table S2 in the Supporting Material shows unevolved and evolved sequences for the protein structures used in this study.

Key quantities measured in simulations are first passage time to the native state, *t*\*, folding stability, *P*_nat_, and native contacts formed. *P*_nat_ is measured as the proportion of steps that the protein is in its native conformation, from the time that the protein reaches the native state until the end of the simulation. Proteins must be exactly in their native conformations to be considered native. For first passage time to the native state, *t* = 0 is defined as the start of the posttranslational phase of simulation in which the protein chain has no conformational restrictions. Native contact counts are either normalized by the maximum possible number of native contacts at a particular chain length or the number of native contacts in the full-length protein (28 for all lattice proteins in this work) and are typically reported as average values for each nascent chain length. For most results in this work, only data for nascent chain lengths of 15 through 27 are reported.

## Results

### Proteins evolve improved stability and kinetics

Beginning with sequences designed for native state thermodynamic stability, we initiated evolutionary simulations. The evolutionary trajectories of the nine proteins are shown across Figs. S3 and S4 in the Supporting Material. In each trajectory, over the course of 1000 mutation attempts, a number of mutations were fixed. Occasionally, deleterious mutations were fixed because fitness evaluations using MC simulations are stochastic. Nonetheless, the fitnesses, folding stabilities, and folding speeds of the evolved sequences are improved over the unevolved sequences. The remaining discussion focuses on the evolved sequences from the end of evolutionary simulation and comparison with unevolved sequences or evolved sequences from alternative evolutionary scenarios.

The overall outcomes of evolutionary simulations are shown in Fig. 2, which compares the fitnesses, folding stabilities, folding times, and native energies of the unevolved, initial sequences to those of the evolved sequences. The total protein activity (Eq. 5), which determines fitness (Eq. 6), is determined by the combination of stability (*P*_nat_) and folding kinetics (the distribution of *t*\*). Here, protein folding stability is shown in terms of the two-state free energy 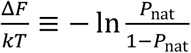 to illustrate differences in folding stability for *P*_nat_ close to 1 more clearly. Since the entire ensemble of first passage times determines fitness, folding time distributions are illustrated using boxplots, with *t* = 0 defined as the moment of release from the ribosome.

**Fig. 2:**
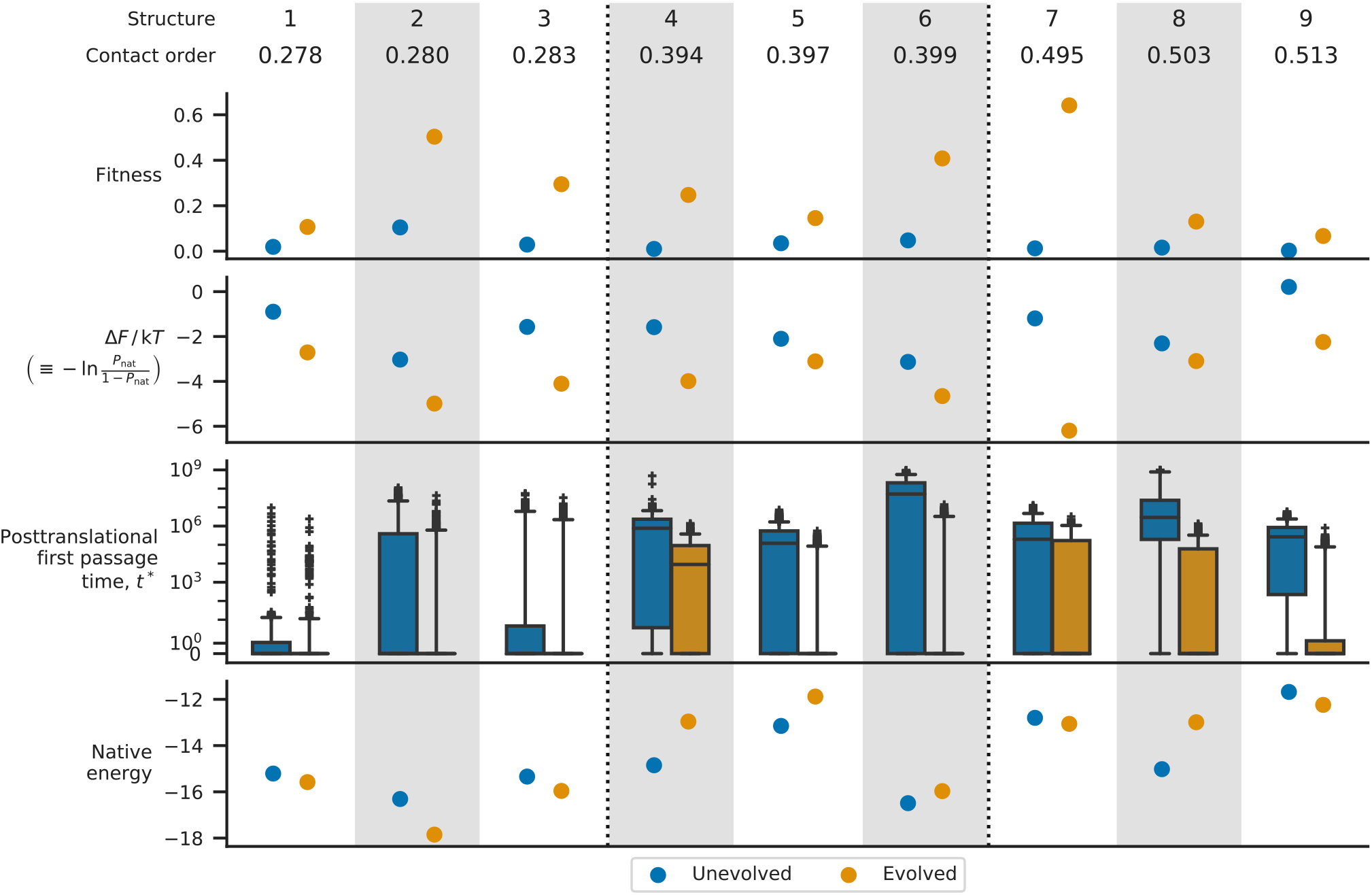
Outcomes of evolutionary simulations. Comparison of properties of initial, unevolved sequences to those of sequences obtained by evolutionary simulation under selection for stability and kinetics (Eqs. 5 and 6). Three groups of three native structures, of low, medium, and high contact orders (numbered 1 through 9, vertical dotted lines indicate grouping), were selected for simulation. The four plots show fitness, folding stability (as 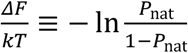), first passage time to the native state, and native state energy for unevolved sequences (blue) and evolved sequences (orange). First passage times are measured using MC simulations of translation and folding, boxplot whiskers show the 5^th^ and 95^th^ percentile values, and *t* = 0 is the moment of release from the ribosome. For unevolved sequences 6 and 8, 4 out of 900 and 27 out of 900 simulations failed to fold, respectively, within the posttranslation period of 10^9^ time units.

Initial, unevolved sequences have moderate stabilities that improve with evolution. The unevolved sequences also fold slowly relative to the degradation timescale (1/*k_d_*) of 200,000 time units: all medium and high contact order unevolved sequences have nonzero median first passage times, meaning that proteins fold posttranslationally in the majority of folding trajectories. The longest first passage times for unevolved sequences are greater than 10^9^ time units, indicating that unevolved sequences enter long-lived kinetic traps. In comparison, evolved sequences fold much faster: only the evolved sequence for structure 4 has a nonzero folding median first passage time, of 9,000 time units. This indicates that cotranslational folding, defined as reaching the native state prior to release from the ribosome, occurs in the majority of folding trajectories for evolved sequences. Since most evolved sequence folding times are well within the degradation timescale, stability becomes the main determinant of fitness for evolved sequences, with evolved sequences folding to structures 2 and 7 having the highest stability and therefore the highest fitnesses. Later sections of this work will examine sequences folding to structures 2 and 7 more closely.

To probe the effect of different selection pressures, evolutionary simulations were also run under two alternative scenarios. The first alternative scenario, “no translation,” simulates protein folding starting from full-length proteins in fully extended conformations. The second alternative scenario, “no folding,” removes the initial first passage to the folded state by starting simulations with proteins already in their native conformations (effectively setting *t*\* in Eq. 5 to 0). The properties of sequences obtained from evolution under these two alternative fitness scenarios are shown in Fig. S5 in the Supporting Material. Note that although evolutionary simulations were performed using alternative fitness evaluations, the fitnesses and first passage times shown in Fig. S5 are obtained from evaluating all sequences using the regular fitness function evaluation which involves simulating protein translation and folding. We observe that evolving for *in vitro* refolding kinetics (“no translation” scenario) results in proteins that fold in the translational context just as fast as our regular sequences which evolved to fold with translation. The evolved “no translation” sequences, except that of structure 5, are less stable than sequences evolved with translation however, suggesting that an ordered build-up of protein native structure during translation allows sequences to evolve higher stability. On the other hand, the results of evolution under the “no folding” scenario shows that long-lived kinetic traps that hamper folding are not eliminated under an evolutionary scenario where folding rates do not affect protein activity and fitness.

The bottom-most plot in Fig. 2 shows the native state energies of unevolved and evolved sequences. We observe that for structures 1, 2, 3, and 6, evolved sequences have low native energies, whereas for structures 4, 5, 7, 8, and 9, evolved sequences have relatively higher native energies. These native energies reflect whether these proteins fold early on or late during translation, as we will discuss next.

### Two opposing strategies for reaching the native state

Our model proteins evolved sufficiently fast kinetics such that the majority of folding trajectories for evolved sequences exhibit cotranslational folding (Fig. 2). To understand the nature of cotranslational folding in our model proteins, we examined the folding trajectories for our unevolved and evolved sequences. We first characterized trajectories by the fraction of possible native contacts formed at each nascent chain length, 〈*Q*〉_*L*_. This measure normalizes the number of native contacts formed at a particular chain length by the maximum number of native contacts that can possibly be formed at that chain length. *Q* is averaged over all trajectory samples at each chain length and over all trajectories, providing an ensemble-average folding trajectory.

In examining 〈*Q*〉_*L*_, we observe that the behavior of our evolved proteins falls into two groups, as illustrated in Fig. 3. Group 1 consists of low contact order structures 1, 2, and 3 as well as medium contact order structure 6. For evolved sequences in Group 1, 〈*Q*〉_*L*_ is close to 1, indicating that nascent chains adopt native-like conformations in which nearly all possible native contacts are formed after growing beyond a chain length of 16 (Fig. 3A). Group 2 consists of medium contact order structures 4 and 5 and high contact order structures 7, 8, and 9. Evolved sequences in Group 2 do not develop high 〈*Q*〉_*L*_ values until nascent chains grow to about 25 residues in length (Fig. 3B). Thus, we characterize Group 1 proteins as folding early on during translation and Group 2 proteins as folding toward the end of translation.

**Fig. 3:**
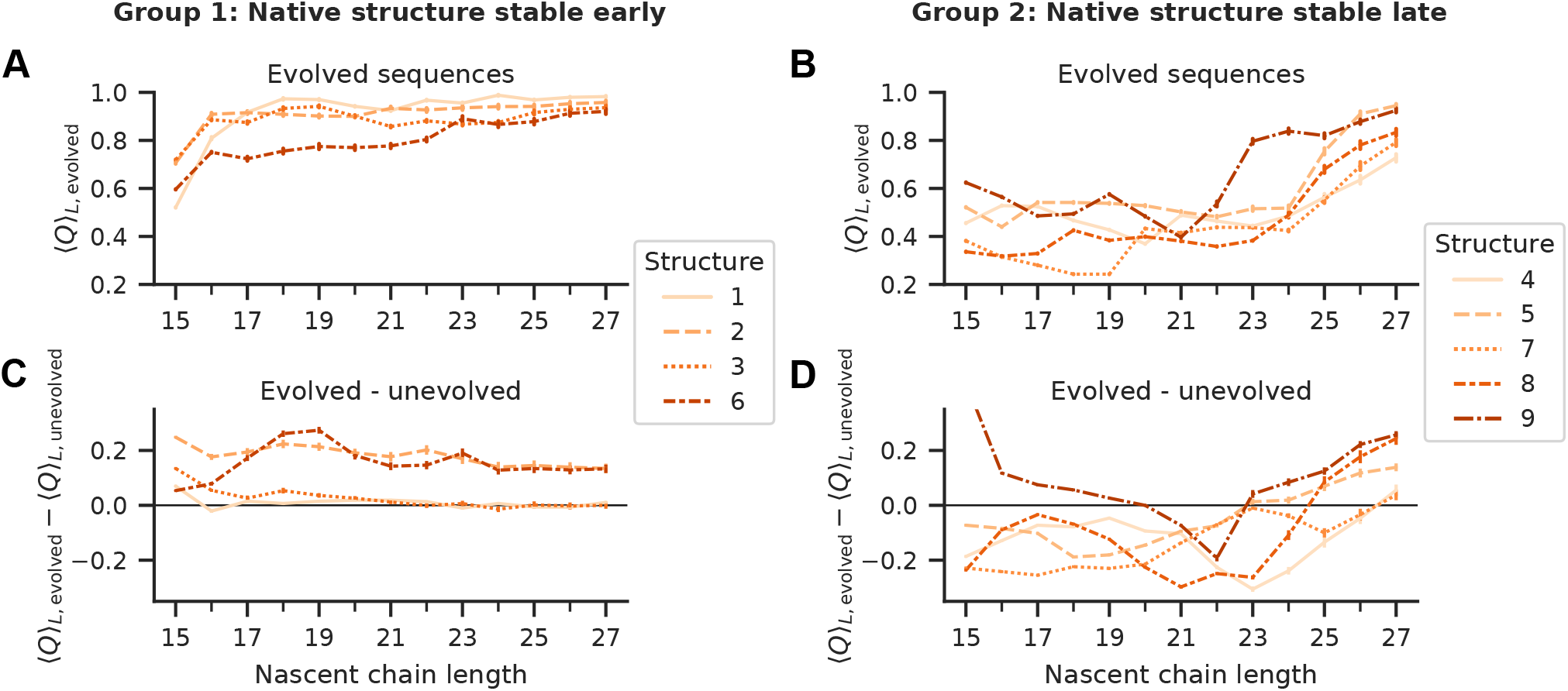
Proteins can be separated into two groups based on behavior during translation. MC simulations of translation and folding for each sequence are illustrated using average *Q* at each chain length, 〈*Q*〉_*L*_. *Q* is the fraction of native contacts formed out of the total possible number of native contacts. Only data for nascent chain lengths 15 through 27 residues are shown here. (A) 〈*Q*〉_*L*_ for evolved protein sequences in Group 1 where the native structure is stable early on during translation. (B) 〈*Q*〉_*L*_ for evolved protein sequences in Group 2 where the native structure is stable toward the end of translation. (C), (D) Same protein structures as in (A) and (B), respectively, but showing difference in 〈*Q*〉_*L*_ between evolved and unevolved sequences. To guide the eye, 0 is indicated by a horizontal line. All error bars indicate 95% confidence intervals obtained by bootstrap sampling per-trajectory values.

There are also differences in how sequences evolved, as illustrated by differences in 〈*Q*〉_*L*_ between evolved and unevolved sequences. For Group 1, 〈*Q*〉_*L*_ either increased or remained unchanged as a result of evolution (Fig. 3C). Minimal change in reflects cases in which 〈*Q*〉_*L*_ is already close to 1 for unevolved sequences (structures 1 and 3). Interestingly, for Group 2, evolved sequences other than the sequence folding to structure 9 have lower 〈*Q*〉_*L*_ values at intermediate nascent chain lengths 15-22 compared to those of unevolved sequences (Fig. 3D) (Mann-Whitney *U* test between per-trajectory values at each length for each sequence, *P* < 0.0001). This indicates that evolved sequences form fewer native contacts at those lengths.

To further understand these two kinds of proteins, we examined individual folding trajectories for each structure. Here, we switch to quantifying folding using native contact counts, to illustrate development of native structure during translation. Native contacts are normalized by the total number of native contacts for the full-length native conformation, which is 28 for all lattice proteins in this work. We focus on structures 2 and 7 as representatives of Group 1 and Group 2, respectively, since evolved sequences for structures 2 and 7 have the highest folding stability out of all nine evolved sequences. Individual folding trajectories for unevolved sequences folding to structures 2 and 7 are shown in Figs. 4A and 4B, respectively, and the corresponding trajectories for evolved sequences are shown in Figs. 4C and 4D, respectively. For each trajectory, native contacts are averaged over all samples collected at each nascent chain length. Folding trajectories for unevolved and evolved sequences of all nine native structures are shown Fig. S6 in the Supporting Material.

**Fig. 4:**
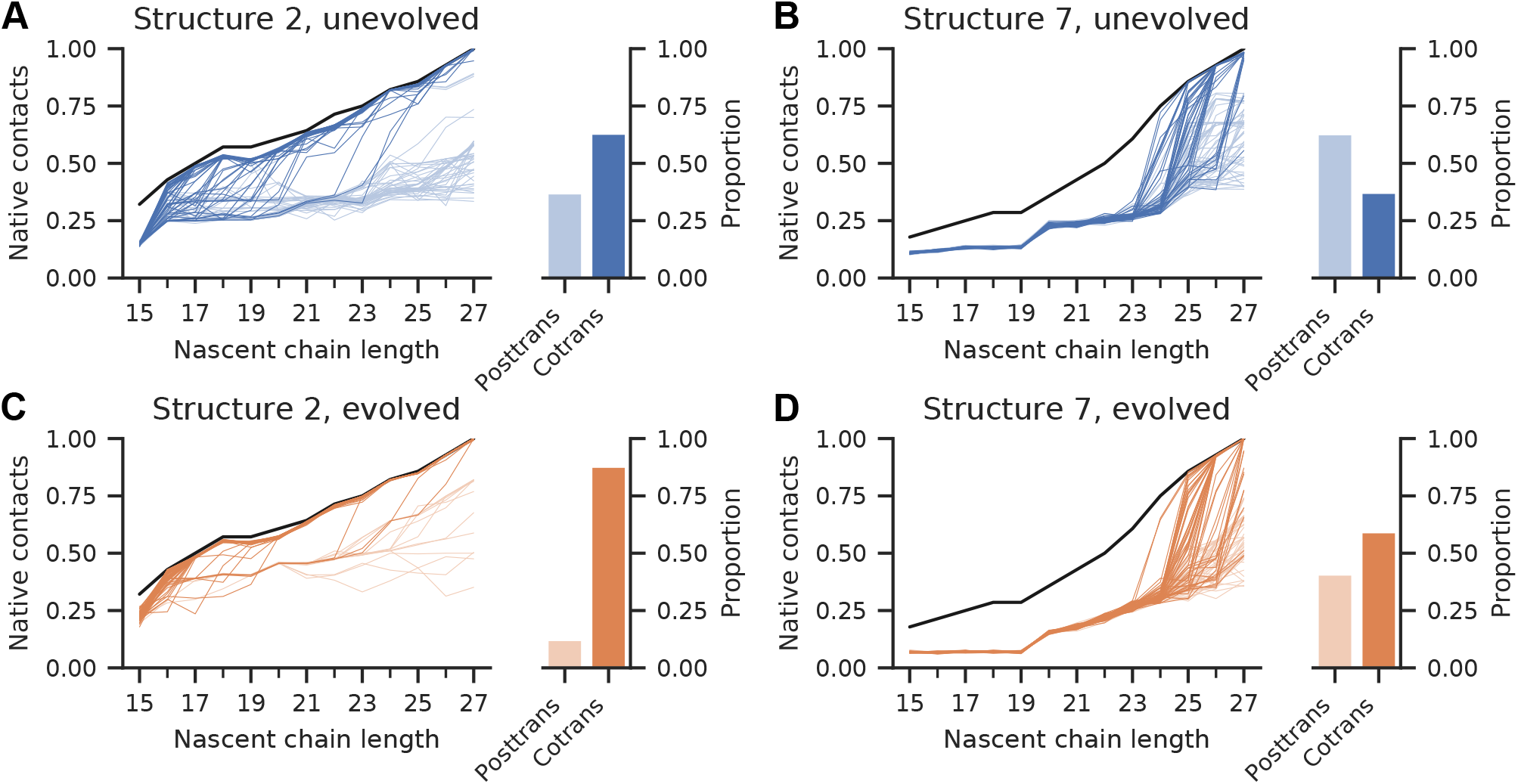
Structures 2 and 7 as respective examples of proteins folding early (Group 1) or late (Group 2) during translation. Individual folding trajectories for sequences folding to structures 2 and 7 are shown by native contacts at nascent chain lengths 15-27. Native contacts are normalized by the total number of native contacts in full-length native structures, 28, and averaged over all samples collected at each nascent chain length. Each colored line is a single trajectory, and each panel shows 100 trajectories. The shade of a colored line or bar indicates whether a particular trajectory folded before (dark) or after (light) release from the ribosome. Solid black line in each panel indicates theoretical maximum number of native contacts at each nascent chain length. (A), (B) Folding trajectories for unevolved sequences folding to structures 2 and 7, respectively. (C), (D) Folding trajectories for evolved sequences folding to structures 2 and 7, respectively. Righthand side of each panel shows proportion of trajectories that reach the native state before (“cotrans”) or after (“posttrans”) release from the ribosome.

Sequences folding to structure 2 can stably occupy native-like conformations beginning at a nascent chain length of 16 residues (Figs. 4A and 4C). There is an apparent bimodality to the folding trajectories, with the majority of trajectories occupying native-like conformations during translation, and a minority of trajectories in partially-native states. Most trajectories reach native-like states when the nascent chain is between 15 and 21 residues in length; trajectories that fail to fold by length 21 mostly fold posttranslationally. For instance, for the evolved sequence, 82% of folding trajectories reach the native state by length 21, but of the remaining trajectories, 67.5% remain unfolded at the end of translation. This demonstrates kinetic partitioning (45); the additionally translated residues produce kinetic traps in the folding energy landscape. The difference between unevolved and evolved sequences is that for the evolved sequence, a greater proportion of trajectories reach the native state while the nascent chain is less than 21 residues in length, resulting in a higher proportion of cotranslational folding. Other proteins in Group 1 are similar, with a majority of trajectories (>75%) released from the ribosome in the native state for evolved sequences (Fig. S6).

In contrast, for sequences folding to structure 7, folding does not occur until the nascent chain reaches a length of 25 residues (Figs. 4B and 4D). Compared to the unevolved sequence, the evolved sequence also forms fewer native contacts at nascent chain lengths below 23 residues. Once the nascent chain passes 25 residues in length however, evolved sequence trajectories show rapid folding to native-like conformations. Other proteins in Group 2 similarly fold only toward the end of translation (Fig. S6). Upon folding, the full native fold, minus just a few contacts, is achieved. Thus, although many evolved sequences in this second group technically demonstrate cotranslational folding (by folding before the end of translation), the folding process is closer to what would occur for full-length proteins.

Contact order is not a perfect predictor of cotranslational folding, as seen by the split of our medium contact order structures among Group 1 and Group 2. In particular, the evolved sequence for medium contact order structure 6 folds early on during translation. To explore this, we constructed 2D contact maps in which average frequencies of residue-residue contacts at particular nascent chain lengths during translation are averaged across all folding trajectories. Figs. S7, S8, and S9 in the Supporting Material show these contact map-based plots of folding trajectories for unevolved and evolved sequences folding to structures 2, 6, and 7, respectively. For sequences folding to structures 2 and 6 (Figs. S7 and S8), evolution strengthened native contacts and weakened nonnative contacts. Both structures 2 and 6 equally support forming 17 native contacts at a nascent chain length of 20. By this point during translation, nascent chains adopt native-like conformations, from which the rest of the native structure forms. These native-like conformations are cotranslational folding intermediates, stable states on the path to the full-length native structure, similar to intermediates we predicted in our previous work (20). Contact order roughly predicts whether such cotranslational folding intermediates can form. In contrast, for structure 7 (Fig. S9), only 10 native contacts can be formed when the nascent chain is 20 residues long, and the nascent chain instead forms more nonnative contacts. For sequences folding to structure 7, evolution weakened both native and nonnative contacts at shorter nascent chain lengths. These observations explain how the evolved sequence for structure 7 forms fewer native contacts at intermediate nascent chain lengths than does the unevolved sequence (Fig. 4B versus Fig. 4D).

We find further distinctions between Group 1 and Group 2 proteins when examining energetics as a function of nascent chain length. The free energies of native conformations are shown in Fig. S10 of the Supporting Material. Mirroring observations made from analyzing native contacts and contact maps, we observe that for Group 1, evolution increased or maintained the stability of native conformations. On the other hand, for Group 2, evolution destabilized native conformations at shorter nascent chain lengths. This destabilization of native conformations is reflected in native state energies as well. Fig. S10 also shows the energies of native conformations for nascent chain lengths 15-27. For proteins in Group 2, we observe that the C-terminal residues for the evolved sequences contribute a greater fraction of the stabilization of the native state than is the case for the unevolved sequences. This pattern explains why evolved native energies for structures folding either early or late during translation are different in magnitude (Fig. 2, bottom). Overall, we find that the native structure of a protein determines how many native contacts are available at a nascent chain length, which decides whether the nascent chain can stably fold. This in turn influences whether evolution strengthens or weakens contacts made by residues at particular nascent chain lengths.

### Kinetic characterizations contrast cotranslational folding with *in vitro* folding

For Group 1 proteins, folding trajectories that do not reach the native conformation while the nascent chain is 15-20 residues long mostly fold posttranslationally (Figs. 4A, 4C, and S6). This suggests that folding kinetics slows with increasing nascent chain length for these proteins. To investigate this, *in vitro* first passage times to the native state—the full-length protein starts in an extended conformation—were measured. We compare first passage times of unevolved and evolved sequences to those of sequences obtained from evolution under the “no translation” scenario (Fig. 5); the latter sequences, by design, are optimized for *in vitro* folding. Compared to the unevolved sequences, both evolved and evolved, “no translation” sequences have faster folding kinetics. More significant, the four evolved sequences in Group 1 (structures 1, 2, 3, and 6) have slower first passage times compared to their “no translation” counterparts as well as compared to sequences in Group 2 (structures 4, 5, 7, 8, and 9). The *in vitro* first passage times of the evolved sequences that fold early on during translation are only moderately improved compared to those of their initial, unevolved counterparts.

**Fig. 5:**
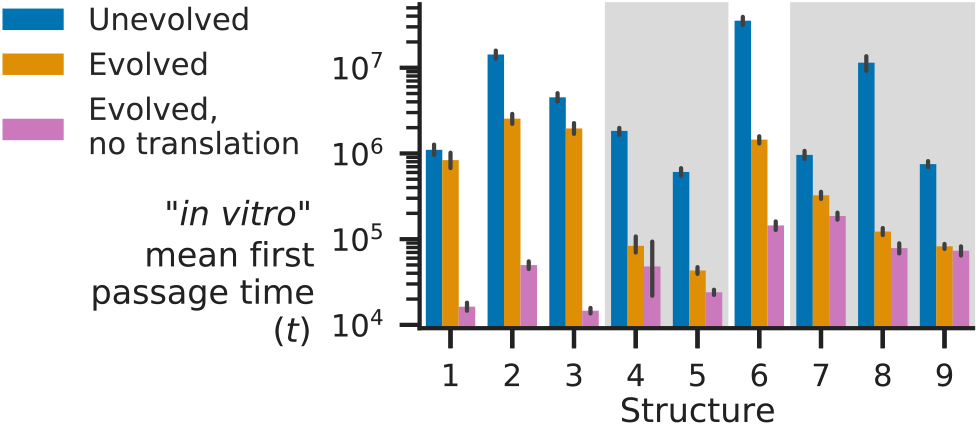
Structures evolved to fold cotranslationally have slow *in vitro* folding kinetics. Mean first passage times for folding from full-length, extended conformations for unevolved sequences (blue), evolved sequences (orange), and sequences evolved in the “no translation” evolutionary scenario (magenta). Structures in Group 2 have been given a shaded background. Error bars indicate 95% confidence intervals obtained by bootstrap sampling.

These differences in kinetics between sequences reflect different selection pressures during evolution. When undergoing translation, evolved sequences in Group 1 fold to stable, native-like conformations at lengths of 15-20 residues. Additional translated residues then add to an existing native structure. Any slow-folding intermediates that form when folding from the fully unfolded state are thereby avoided. One consequence demonstrated here is that proteins that have evolved to fold cotranslationally have slow *in vitro* folding kinetics. Vectorial synthesis reduces the selection pressure for fast folding kinetics when proteins can start folding cotranslationally.

We further characterized the kinetics of our sequences by measuring first passage times to native conformations at chain lengths of 15 through 27 residues. We fit our data to a simple three-state, three-parameter model (see Extended Methods and Fig. S1 of the Supporting Material). The fitted kinetic parameters are shown in Fig. S11 of the Supporting Material. These results show how proteins in Group 1 have fast folding kinetics at intermediate chain lengths and slower kinetics as chain length increases.

### Proteins that fold early on during translation benefit from mid-sequence slow codons

Thus far, our studies have examined nonsynonymous sequence changes, but another aspect of protein translation is that codon identity influences translation rates, which can affect protein folding efficiency (46). Furthermore, slowly translating rare codons represent a bioinformatic signature of cotranslational folding (19, 20). We next investigated the effects of changing the elongation intervals for different codon positions. Here, we restrict our investigation only to evolved sequences.

Estimated translation rates for different codons in *E. coli* differ by up to an order of magnitude (47). We simulated the effect of rare codon substitution on translation speeds by increasing the elongation interval of individual residues tenfold (from 20,000 to 200,000 time units). Note that increasing the elongation interval for the Nth codon means that the translating protein spends additional time while at a length of N-1 residues. We performed MC simulations of translation and folding, increasing the elongation interval for single residues and measuring the proportion of trajectories folding cotranslationally. Results from these simulations for evolved sequences folding to structures 2 and 7 are shown in Fig. 6A and Fig. *6*B, respectively, and results for all evolved sequences are available in Fig. S12 of the Supporting Material. Introducing slowly translated codons can substantially increase the proportion of folding trajectories folding cotranslationally. For nearly all sequences, increasing the elongation interval of codons at the C-terminus increases the proportion of cotranslational folding (Fig. S12), with proteins in Group 2 showing more substantial increases in cotranslational folding. Increased cotranslational folding due to C-terminal slow codons is a somewhat trivial effect under our model however, since folding at a nascent chain length of 25 or 26 residues is not very different from folding as a full-length 27-residue protein. Only sequences folding to structures 2 and 6, from Group 1, show increases in cotranslational folding from increased elongation intervals at mid-sequence positions. These positions reflect nascent chain lengths at which cotranslational folding intermediates become stable, and resembles how rare codons are positioned before putative cotranslational folding intermediates in real proteins (20).

**Fig. 6:**
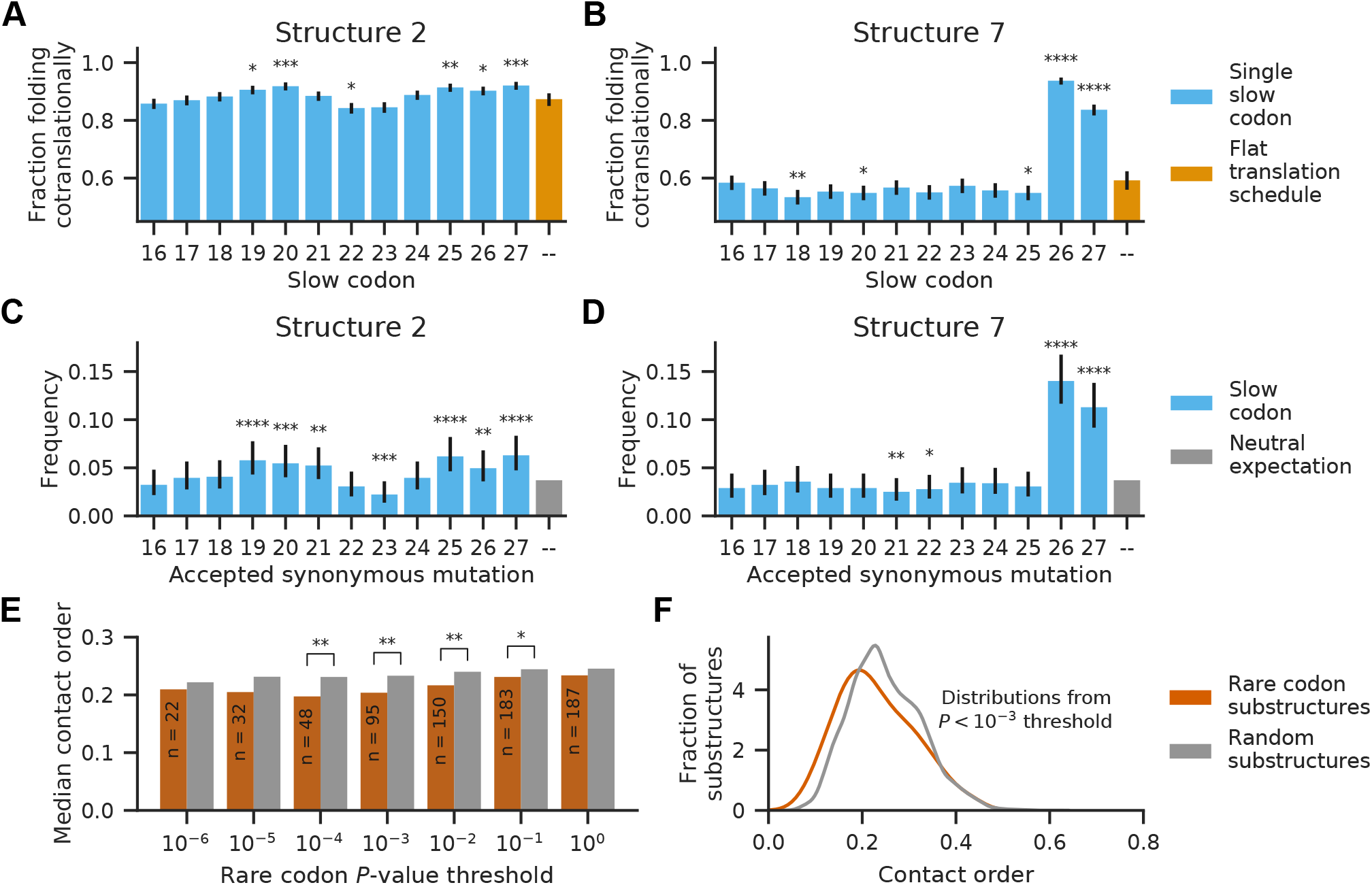
Slowing translation at particular mid-sequence positions enhances cotranslational folding in lower contact order proteins. (A), (B) Proportion of folding trajectories with cotranslational folding when translation is slowed at single codon positions for evolved sequences folding to structures 2 and 7, respectively. 1500 folding simulations were performed for each slow codon position, and 900 folding simulations were performed for the original, flat translation schedule. Error bars indicate 95% confidence intervals calculated by Wilson score interval. Statistical significances of differences in fraction folding cotranslationally (compared to translation using a flat translation schedule) were evaluated using chi-squared tests. (C), (D) The frequency that a synonymous mutation was fixed in synonymous mutation evolutionary simulations for evolved sequences folding to structures 2 and 7, respectively. Frequencies for positions 1-15 are omitted. 1800 independent evolutionary simulations were performed for each sequence. Error bars indicate 95% confidence intervals calculated using Goodman’s method. Statistical significances of deviations from the neutral expectation of 1/27 were evaluated using independent binomial tests. (E) The median contact order of protein substructures preceding evolutionarily conserved rare codons compared to the median contact order of substructures preceding random positions in genes without rare codons at different *P*-value thresholds for evolutionary conservation of the rare codons. The number of genes with conserved rare codons at each *P*-value threshold is indicated inside the bars; the remaining genes without rare codons were used to generate random substructures with lengths distributed according to a geometric distribution. Distributions of contact orders were tested for statistically significant differences using the Mann-Whitney *U* test (two-sided). (F) The distributions of contact orders for substructures preceding rare codons and for substructures preceding random positions at the 10^−3^ *P*-value threshold for evolutionary conservation. For all panels, *: *P* < 0.05, **: *P* < 0.01, ***: *P* < 0.001, ****: *P* < 0.0001.

To confirm that increases in cotranslational folding proportion could provide a fitness advantage and therefore be selected by evolution, we performed evolutionary simulations in which mutations were restricted to slowing translation at specific positions in the sequence, mimicking the effect of synonymous mutation to a rare codons. In these evolutionary simulations, simulation trajectories were stopped once a single synonymous mutation was fixed (see Extended Methods). The distribution of fixed slow codon substitutions for evolved sequences folding to structures 2 and 7 are shown in Fig. 6C and Fig. 6D, respectively; the distributions deviate significantly from the neutral expectation of a uniform distribution (chi-squared test, *P* < 0.0001). As predicted by the cotranslational folding proportions in Fig. 6A and Fig. 6B, synonymous substitutions to slower codons are selected for by our evolutionary simulations.

Our model results suggest that proteins that can fold early on during translation benefit from translational slowing at specific mid-sequence positions. Since our model proteins that fold early on during translation are also lower in contact order, we wondered whether a bioinformatic signature of cotranslational folding, conserved rare codons, would be more likely to occur for proteins with lower contact orders. A recent study from our group identified conserved rare codons in *E. coli* and found that such rare codons are frequently positioned downstream of cotranslational folding intermediates, which were independently identified according to a native-centric folding model (20). We used the rare codons identified in this previous study to measure the contact orders of protein substructures. Here, a substructure is defined as the portion of the native structure from the N-terminus to 30 residues before the location of a codon of interest, which accounts for the ribosome exit tunnel (48).

For each gene with rare codons, we looked for the first evolutionarily conserved rare codon and measured the contact order of the substructure corresponding to that codon. We then compared this distribution of contact orders to control distributions generated by measuring the contact orders of substructures preceding random positions in genes without conserved rare codons. This analysis was performed at multiple *P*-value thresholds for determining evolutionary conservation of rare codons (see Extended Methods). We found that protein substructures preceding rare codons have lower contact orders than those of protein substructures preceding randomly drawn positions (Fig. 6E). Statistical significance declines with decreasing *P*-value threshold as the number of genes with qualifying rare codon regions decreases. The most statistically significant difference is found at a *P*-value threshold for rare codon conservation of 10^−3^ (Mann-Whitney *U* test (two-sided), *P* = 0.0014). The distributions at this conservation threshold are shown in Fig. 6F and have medians of 0.2038 and 0.2332 for rare codon substructures and random substructures, respectively. The same analysis was performed using long-range order, an alternative measure of distant residue contacts (49), and even greater differences between distributions of long-range order were observed (Fig. S13 in the Supporting Material).

## Discussion

Our simulations use a simplified model of protein translation and folding to study how sequences evolve under selection pressure to avoid degradation. The fitness function in our evolutionary model was conceived by considering that proteins can begin to fold during translation and that proteins are vulnerable to degradation while free in solution when not in their native states. We further assume that the folded native state is the functional state for our proteins. In our model, the fitness of a protein sequence depends on its native state stability and folding rate. Since the topology of a protein’s native fold affects which native contacts are accessible to nascent chains emerging from the ribosome, we simulated evolution of model proteins with a range of contact orders. We find that whether native-like states are stable early on during translation shapes what folding pathways proteins evolve to optimize.

Our results are summarized in Table 1. When a protein can fold early on during translation, the protein folds cotranslationally by first folding to a partial-length native structure in which all possible native contacts are formed. Subsequent translated residues then add additional native contacts to this core structure. Evolution strengthens native contacts and weakens nonnative contacts to stabilize the native state. Four of our nine proteins—the three low contact order proteins and one medium contact order protein—follow this pattern. As structural analyses of our proteins found, contact order is a rough indicator for whether the protein native structure supports stable, partial-length conformations that facilitate cotranslational folding. One consequence of evolution in this direction is that there is less selection pressure on the folding kinetics of the full-length chain, since most folding trajectories involve residue-by-residue cotranslational folding. We observe that for our proteins that fold early on during translation, *in vitro* folding times—measured from full-length extended conformations—are substantially longer than cotranslational folding times. Experiments on individual proteins have found that refolding from the denatured state is often less efficient than cotranslational folding in terms of folding rate or occurrence of irreversible aggregation (6, 15, 17, 18). Our simulation results suggest one evolutionary factor for these phenomena: proteins that fold cotranslationally are not under selection to avoid forming slow-folding intermediates encountered when refolding from the denatured state. Consequently, we predict that proteins that fold cotranslationally are more prone to inefficient refolding from denatured states.

**Table 1:** Comparison between proteins that fold early or late during translation.

| **Folding early on during translation** | **Folding toward the end of translation** |
| --- | --- |
| Lower contact order | Higher contact order |
| Evolution strengthens contacts | Evolution weakens nascent chain contacts |
| Folds cotranslationally | Folds posttranslationally |
| Full-length protein may have slow folding kinetics | Folding pathways on and off the ribosome are likely to be similar |
| Rare codons mid-sequence or at the C-terminus | Rare codons at the C-terminus |

Interestingly, our observation of slow *in vitro* folding kinetics for our evolved proteins which fold early on during translation contradicts the expected relationship between contact order and folding speed (28). This difference may be because the study of contact order and folding speed has been limited to small proteins capable of *in vitro* refolding (28, 50–53). Indeed, many proteins are unable to refold once denatured *in vitro* (54–58). Although the model proteins in this study are admittedly short in length, their properties can still generalize onto the characteristics of longer, real proteins. We speculate that fast folding kinetics for partial-length nascent chains and slow full-length folding kinetics provides cells with a route for efficient production of long-lived, kinetically stable proteins which, once folded on the ribosome, remain protected from transient unfolding by a high folding-unfolding barrier.

On the other hand, if native-like states are not stable until the protein is nearly fully translated, protein sequences evolve so that the nascent protein chain avoids making native and nonnative contacts that are too strong and trap the nascent chain in incompletely folded states. This pattern of evolution occurs for five of our higher contact order proteins. We observe that compared to unevolved, initial sequences, evolution weakens both native and nonnative contacts formed by the N-terminal portion of the protein. Once a sufficient number of residues are extruded, rapid folding to the native state commences. Although it would be difficult to test whether protein sequences are optimized to avoid forming strong inter-residue interactions unless native-like conformations are stable, it is known that cotranslational chaperones such as trigger factor prevent nascent chains from making aberrant interactions and alter folding pathways (24, 59, 60).

Our model proteins that fold toward the end of translation are higher contact order proteins. While their folding kinetics are sufficiently fast to fold prior to the end of translation in our simulations, the folding pathways of such proteins while tethered are not likely to differ from *in vitro* folding pathways. Recent studies on two proteins, the Src SH3 domain and titin I27, observed that ribosome-nascent chain complex folding pathways are similar to off-ribosome folding pathways (61, 62). We calculate the contact order of these two proteins to be 0.37 and 0.41, respectively; these values are much higher than the median contact order of the *E. coli* proteins used in our bioinformatics analysis, 0.21. Our simulation results predict that such high contact order proteins should fold toward the end of translation or posttranslationally, which agrees with the experimental findings.

We also investigated the effect of changing the elongation interval for specific positions along our evolved sequences to simulate the effect of substitution to rare, slowly translating synonymous codons. These results show that slowly translating codons increase the folding efficiency of our proteins and provide an example of evolutionary selection on synonymous codons. Our results are in agreement with a recent study that showed that synonymous substitutions in a gene diminish fitness by increasing protein degradation (63). Increasing the elongation interval at mid-sequence positions increases folding efficiency only for model proteins that can fold early on during the translation process. We used an existing dataset of conserved rare codons in *E. coli* genes to probe whether contact order has any association with conserved rare codons (20). By comparing the contact orders of protein substructures preceding conserved rare codons to the contact orders of substructures preceding random positions in genes without conserved rare codons, we find that the contact orders of the former distribution is lower than the contact order of the latter. Our findings show that native structure topology indeed influences whether a nascent chain is likely to cotranslationally fold, with protein substructures with more local topologies (and lower contact order) more likely to precede rare codon stretches in real genes.

In conclusion, our simulations have found that the point at which native-like conformations become thermodynamically stable during translation influences how proteins evolve to fold during translation. Our work predicts that cotranslational folding is more likely to occur in lower contact order proteins. Overall, we find that cotranslational folding is a major factor in evolutionary selection of functional proteins that avoid intracellular degradation.

## Author Contributions

Designed research, W.M.J., E.I.S., and V.Z.; Developed theoretical models, W.M.J., E.I.S., and V.Z.; Performed research, V.Z; Analyzed data, E.I.S. and V.Z. Wrote the manuscript, E.I.S. and V.Z.

## Acknowledgments

All computations in this work were run on the FASRC Odyssey and Cannon clusters supported by the FAS Division of Science Research Computing Group at Harvard University. Lattice protein renderings were produced using Tachyon (64) within VMD (65). We thank Rostam M. Razban and Mobolaji Williams for helpful discussions. This work was supported by RO1 GM068670 from NIGMS and a National Science Foundation Graduate Research Fellowship (awarded to V.Z.).

## Supporting citations

References (66–73) appear in the Supporting Material.

